# Adversarial childhood events are associated with Sudden Infant Death Syndrome (SIDS): an ecological study

**DOI:** 10.1101/339465

**Authors:** Eran Elhaik

## Abstract

Sudden Infant Death Syndrome (SIDS) is the most common cause of postneonatal infant death. The *allostatic load hypothesis* posits that SIDS is the result of perinatal cumulative painful, stressful, or traumatic exposures that tax neonatal regulatory systems. To test it, we explored the relationships between SIDS and two common stressors, male neonatal circumcision (MNC) and prematurity, using latitudinal data from 15 countries and over 40 US states during the years 1999-2016. We used linear regression analyses and likelihood ratio tests to calculate the association between SIDS and the stressors. SIDS prevalence was significantly and positively correlated with MNC and prematurity rates. MNC explained 14.2% of the variability of SIDS’s male bias in the US, reminiscent of the Jewish myth of Lilith, the killer of infant males. Combined, the stressors increased the likelihood of SIDS. Ecological analyses are useful to generate hypotheses but cannot provide strong evidence of causality. Biological plausibility is provided by a growing body of experimental and clinical evidence linking adversary preterm and early-life events with SIDS. Together with historical evidence, our findings emphasize the necessity of cohort studies that consider these environmental stressors with the aim of improving the identification of at-risk infants and reducing infant mortality.

## Introduction

Sudden Infant Death Syndrome (SIDS) occurs when a seemingly healthy infant (0-12 months) dies unexpectedly in sleep with no apparent cause of death being established [1]. SIDS is the leading cause of infant death in many developed countries [2], accounting for ~2,700 deaths annually in the US [3]. As such, SIDS has received much attention in the literature [e.g., 4].

The *allostatic load hypothesis* for SIDS [5] purports that prolonged and repetitive exposure to stressors (e.g., poor postnatal weight gain [6], hyperthermia [7], and maternal smoking [8]) is maladaptive and has a cumulative effect that increases the risk of SIDS. It differs from the *triple risk hypothesis* [5], which posits that SIDS is caused by an exposure to intrinsic and external factors during a critical developmental stage. That hypothesis cannot explain the four main characteristics of SIDS, namely male predominance (60:40) in the US, the 39% lower SIDS rate among US Hispanics compared to non-Hispanics [9], the seasonal variation with most cases occurring in winter [10], and that 50% of cases occur between 7.6 and 17.6 weeks after birth with only 10% past 24.7 weeks.

To test the predictions of the *allostatic load hypothesis* for SIDS, we identified two common stressors [5], male neonatal circumcision (MNC) and premature birth, for which latitudinal data were available and tested their association with SIDS. Both stressor are male-biased [11] and may explain the male predominance of SIDS, whereas the first stressor may also explain the lower SIDS rates in Hispanics. MNC is associated with intraoperative and postoperative risks including bleeding, shock, sepsis, circulatory shock, and hemorrhage [12–14] that can result in death [14, 15]. MNC may also cause severe and long-lasting pain, trauma, and psychological impairment due to the circumcision procedure that involves maternal separation, restraint to a board, and the removal of sensitive penile tissues that contain numerous nerve endings [16–21]. Since MNC preference is largely cultural, populations can be classified into Anglophone countries (high MNC rate) and non-Anglophone countries (medium to low MNC rate [22, 23]) (Table S1). If MNC is a risk factor for SIDS, SIDS rates would be higher in Anglophone countries, where MNC is highly prevalent [22], compared to non-Anglophone countries, which traditionally have opposed circumcision [23]. US populations also differ in their MNC practices. Between 2005 and 2010, non-Hispanic Whites were the largest group performing MNC (90.8%), followed by non-Hispanic Blacks (75.7%) and Mexican Americans (44%). If MNC is a risk factor for SIDS, in addition to their low SIDS rates we can also expect Hispanic populations to exhibit lower male bias than non-Hispanics.

Prematurity (birth at a gestational age of less than 37 weeks) is a known risk factor for SIDS [24, 25]. The risk factors unique to preterm infants likely have multiple etiologies and include biological vulnerabilities and prolonged exposure to multiple stressors during the hospitalization in the neonatal intensive care unit (NICU) [24], which elevates the allostatic load and the risk for SIDS [26].

We tested the association of SIDS with these two stressors using prevalence data from 16 worldwide populations, 15 countries, and over 40 US states. This is the first ecological link study to test the association between SIDS and MNC, and it is done at a time that SIDS rates remain alarmingly high [27] two decades after the “Back to Sleep” (BTS) campaign.

## Methods

### Data collection

Global male neonatal circumcision (MNC) rates per country (2005-2006) were obtained by searching for ‘neonatal circumcision’ and country in Google Scholar, Google, and PubMed. Similarly to [28], MNC rates for the remaining countries that could not be obtained through peer reviewed journals and whose adult circumcision rates were estimated by the WHO to be <20% [22] were estimated from the total percentage of Muslims [29] and Jews [30] in the country, as both populations were reported to have 100% circumcision rate [31]. US statewise MNC rates (2009-2013) were obtained from [32].

We collected global SIDS prevalence data for 15 countries from 2004-2013 [9, 33, 34]. Year-matched MNC and SIDS data were available for Australia, Canada, New Zealand, and US. Global SIDS and MNC data are summarized in Table S1. All US SIDS data were obtained from the Centers for Disease Control and Prevention (CDC) Wonder [9]. US statewise MNC rate and male:female SIDS ratio are summarized in Table S2. US statewise male bias SIDS data (1000*M_SIDS_ rate/F_SIDS_ rate) between 1999 and 2016 are summarized in Table S3.

US population data were obtained from the US Census (2000, 2010) [35] and the 2012-2016 American Community Survey 5-Year Estimates [36] (Table S4).

Global prematurity data (2006, 2010) were obtained from the March of Dimes Foundation [37] (Table S5). US statewise prematurity data were obtained from the CDC [38] (Table S6).

Finally, we assembled a global best-year matched MNC (Table S1), SIDS (Table S1), and prematurity rates from various sources (Table S7) for the likelihood ratio test. Likewise, US statewise, best-year matching MNC (Table S2), SIDS [9], and prematurity data [39] were collated (Table S8).

### Data analyses

The global SIDS prevalence map was plotted with R package rworldmap [40]. All correlations were calculated using Pearson correlation. Linear regression analyses performed using ‘lm’ function. Likelihood ratio tests were performed using the R package ‘epicalc’ [41]. Analyses were done in R v.3.4.1. All data and code used in our analyses are available at GitHub (https://github.com/eelhaik/SIDS_eco_study).

## Results

### SIDS prevalence

SIDS prevalence varied greatly among the studied countries, ranging from 0.06 to 0.82 per 1,000 births (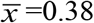, *σ*=0.22) (Figure 1). SIDS prevalence was the lowest in the Netherlands (0.06) and highest in the US (0.82) and New Zealand (0.8). The average SIDS prevalence in the US was notably high compared with Europe (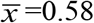, *σ*=0.37) New York had the lowest SIDS prevalence (0.14) and Montana the highest (1.64).

**Figure 1.**
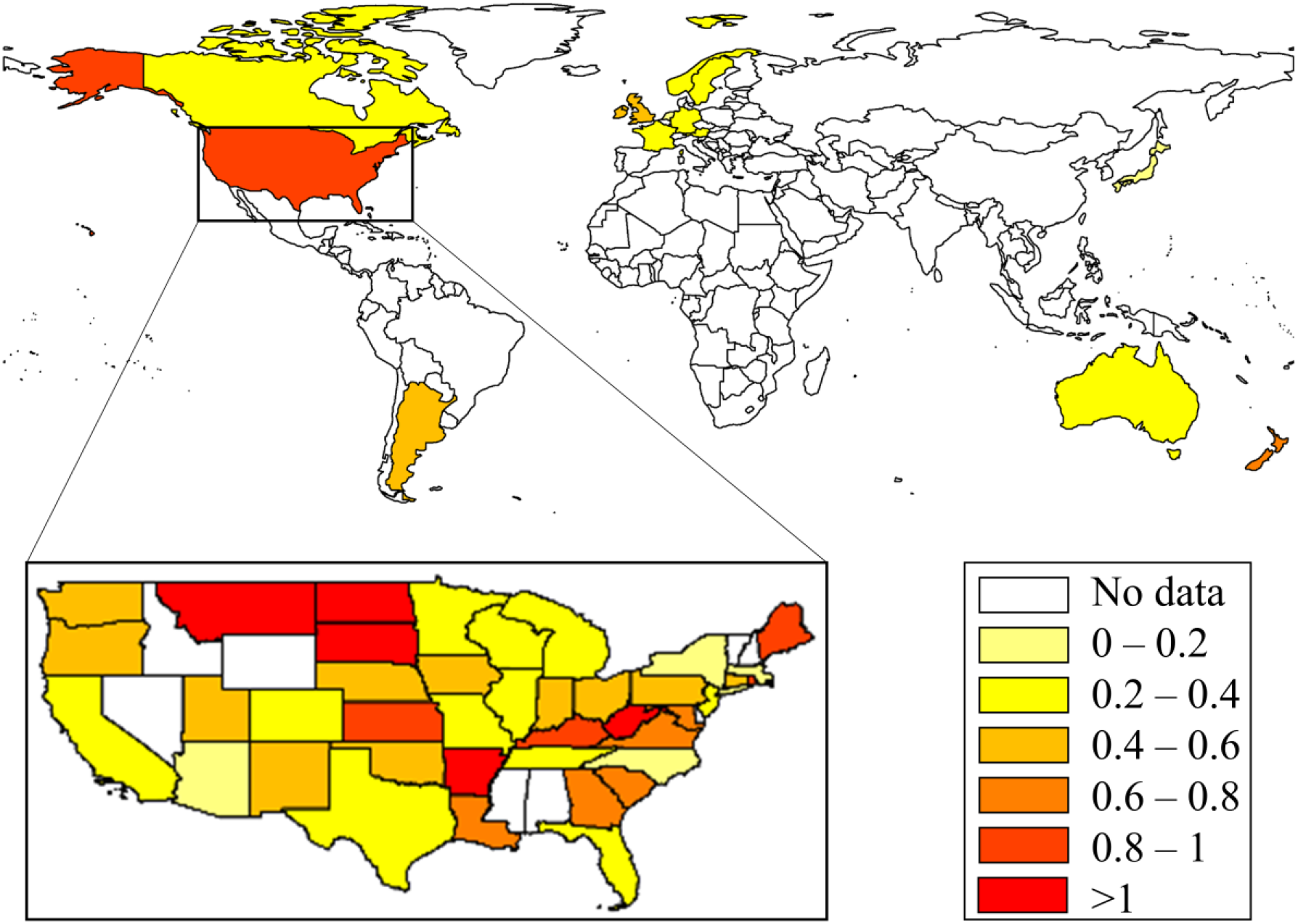
SIDS prevalence (per 1000 births) in 15 countries and 40 US states (inset). SIDS prevalence is color-coded.

Considering the median proportion of US Hispanic (8.4; 2000-2015) in our dataset as a cutoff, the US SIDS prevalence was significantly lower in states (*Nh*) with high Hispanic population (>8% of total population) than state with low Hispanic population (*N_l_*) (Student’s *t*-test 2000: *N_h_*=13, *N_l_*=29, *p-value*=0.047; Δ_SIDS_=0.2; 2010: *N_h_*=25, *N*_l_=20, *p-value*=0.009; Δ_SIDS_=0.28; 2012: *N_h_*=23, *N_l_*=19, *p-value*=0.008; Δ_SIDS_=0.3; 2015: *N_h_*=24, *N_l_*=19, *p-value*=0.002; Δ_SIDS_=0.36). US SIDS prevalence was also significantly negatively correlated with the percent of Hispanics in the population each year (Student’s *t*-test 2000: *N*=44, *r*=-0.37, *p-value*=0.01; 2010: *N*=46, *r*=-0.43, *p-value*=0.003; 2012: *N*=44, *r*=-0.43, *p-value*=0.005; 2015: *N*=44, *r*=-0.41, *p-value*=0.007) (Figure 2, Table S4).

**Figure 2.**
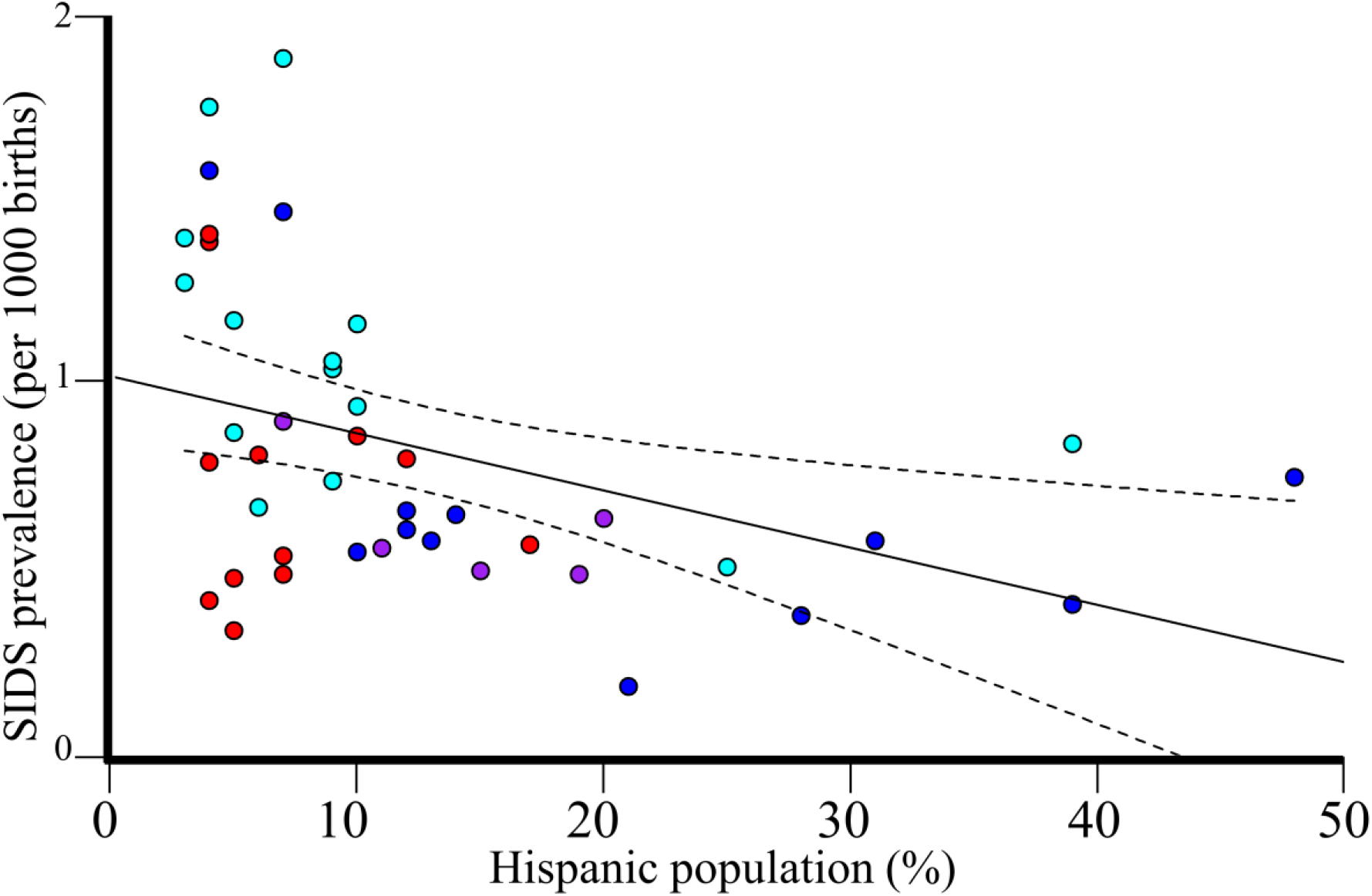
Regression analysis of Hispanic population in the US and SIDS prevalence in 2016. The 95% confidence intervals of the best fit line are denoted in dashed lines. Colors correspond to four US regions: Northeast (violet), Midwest (red), South (cyan), and West (blue).

### MNC is positively associated with the risk for SIDS

The global SIDS prevalence and MNC rates are strongly and significantly correlated (*N*=16, *r*=0.7, Student’s *t*-test, *p-value*=0.003) (Figure 3). The results remain significant even if the MNC rates for the estimated cohort are halved or doubled (*r*=0.69, *p-value*=0.003, 95% CI: 0.004-0.015). The slope of this trend indicates that a 10% increase in the MNC rates is associated with an increase of 0.1 per 1000 SIDS cases. Anglophone countries practice significantly more MNC and have significantly higher SIDS prevalence than non-Anglophones (Kolmogorov-Smirnov test; *p-value*=0.017 and *p-value*=0.007, respectively) (Figure 4).

**Figure 3.**
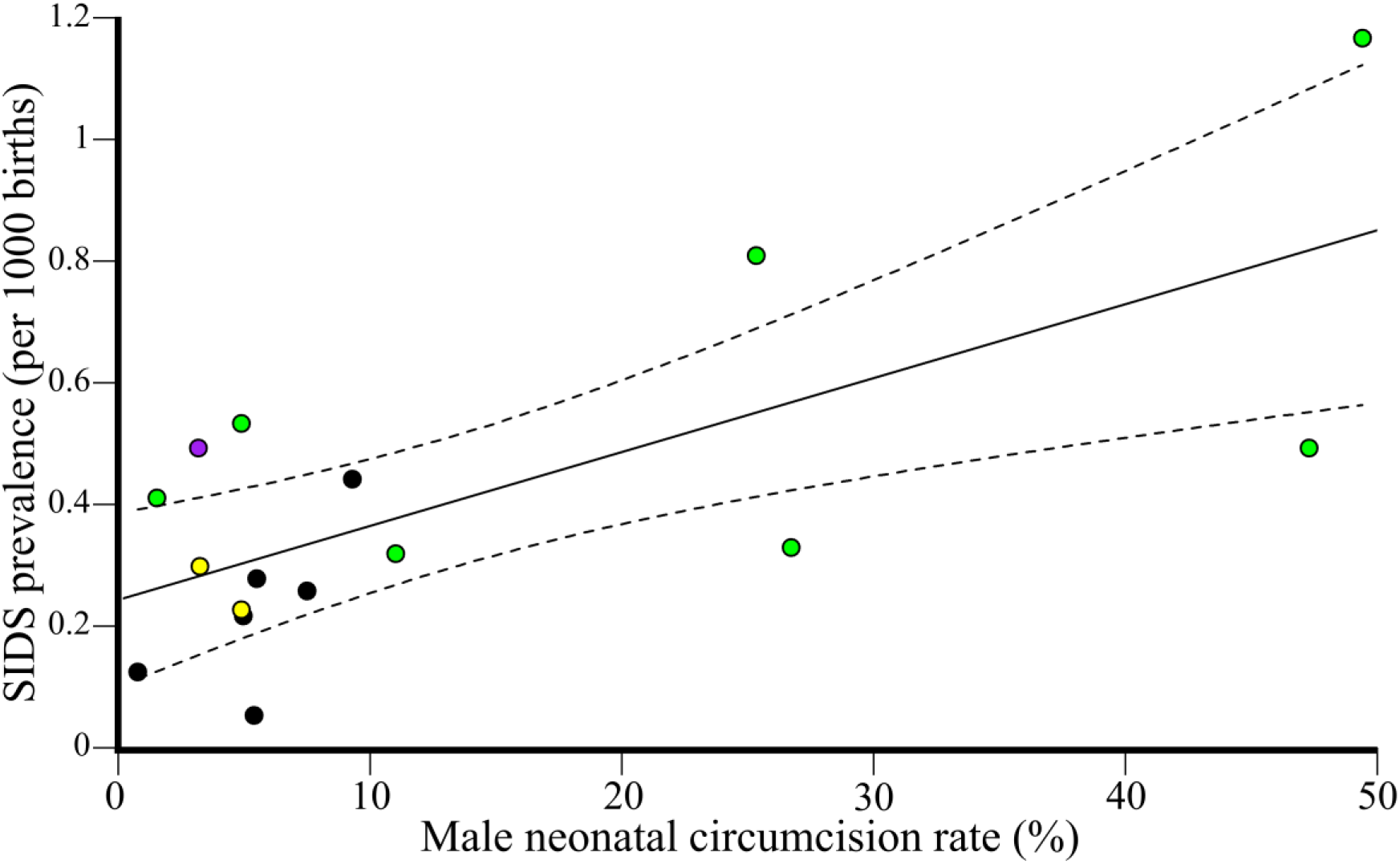
Regression analysis of global MNC rate and SIDS prevalence. The 95% confidence intervals of the best fit line are denoted in dashed lines. Colors correspond to the four population groups: Anglophone countries (green), Ibero-American countries (violet), Nordic countries (yellow), and all other (black).

**Figure 4.**
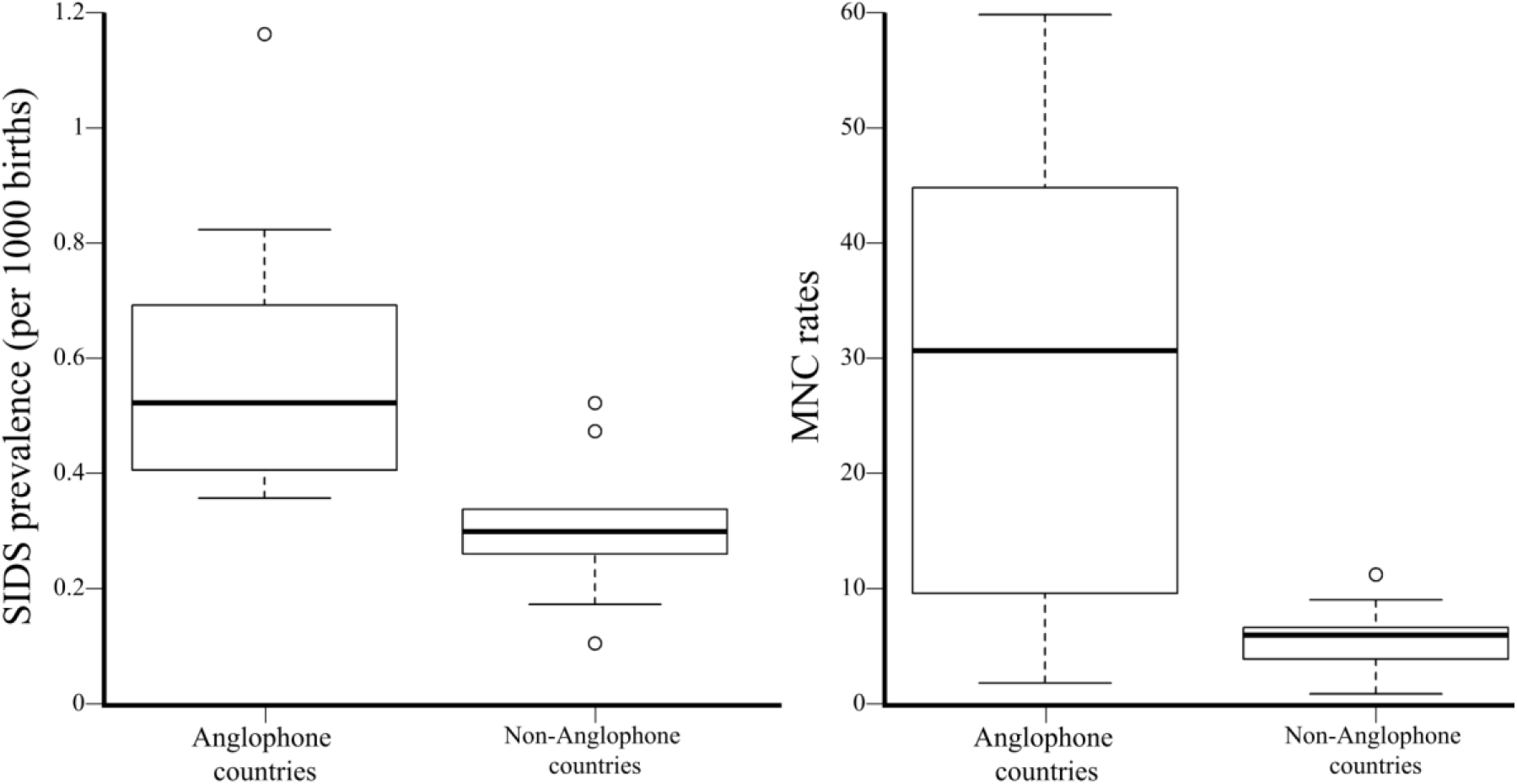
A comparison of the SIDS prevalence and MNC rate in 8 Anglophone and 10 non-Anglophone countries using boxplots.

The US statewise SIDS prevalence and MNC rates are strongly and significantly correlated (*N*=39, *r*=0.37, Student’s t-test, *p-value*=0.02, 95% CI: 0.06-0.61) (Table S8). The slope of this trend indicates that a 10% increase in the MNC rates is associated with an increase of 0.05 per 1000 SIDS cases. In 22 out of 33 states where Medicaid, the most common US health insurance, covers MNC, the average MNC rate (72.4) is nearly twice than the MNC rate in other states (34.3), the SIDS prevalence is marginally significantly higher (0.77 vs 0.5, Kolmogorov-Smirnov test, *p-value*=0.07), and the male bias is higher (1.52 vs 1.23, Kolmogorov-Smirnov test, *p-value*=0.13), but not significantly so (Figure 5).

**Figure 5.**
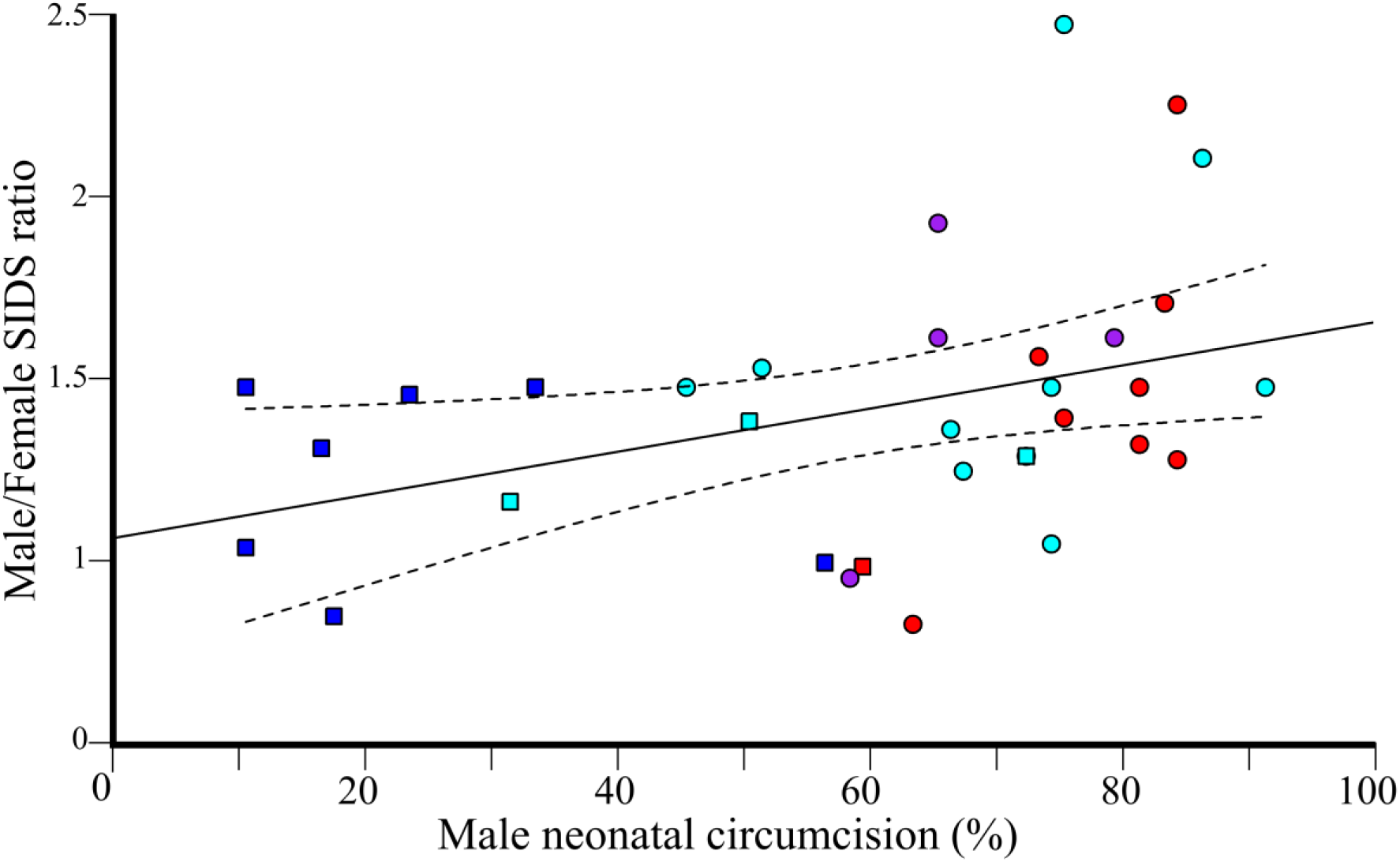
The contribution of MNC toward SIDS gender bias in the US. Symbols mark states where Medicaid, the leading insurance company in US, covers (squares) or does not cover (circles) MNC. The 95% confidence intervals of the best fit line are denoted in dashed lines. Color codes are as in Figure 2.

To test whether the gender bias in SIDS can be explained by MNC, we compared the MNC rates and male:female SIDS ratios in non-Hispanics Whites and Blacks (Table S2). There is a significantly high correlation between the MNC rate and SIDS gender ratio (*N*=33; *r*=0.38, Student’s *t*-test, *p*-value=0.03, 95% CI: 0.04-0.64) (Figure 5). MNC explains 14.2% of the variability in male SIDS deaths in the US. Moreover, of the three largest population groups in the US (Table S3), non-Hispanic Whites (NHW) have the highest MNC [42]. NHW also have significantly higher SIDS male bias compared to Hispanic Whites (HW) (*N*=18, Kolmogorov-Smirnov test, NHW=1.42, HW=1.35, *p-value=0.04)* and non-Hispanic Blacks (NHB) (*N*=18, Kolmogorov-Smirnov test, NHW=1.42, BNH=1.28, *p-value*=6.11*10^−5^).

### Prematurity is positively associated with the risk for SIDS

To test the association between prematurity and SIDS, we considered the global and US prematurity rates. Prematurity rates (%) are the highest in the US (18.46% and 11.7% for non-Hispanic Blacks and Whites, respectively) and lowest in Nordic and East Asian countries (Table S5). The global SIDS prevalence and prematurity rate are strongly and significantly correlated (*N*=16, *r*=0.7, Student’s *t*-test, p-value=0.002, 95% CI: 0.31-0.89) (Figure 6). The slope of this trend indicates that a 10% increase in the prematurity rates is associated with an increase of 0.6 per 1000 SIDS cases.

**Figure 6.**
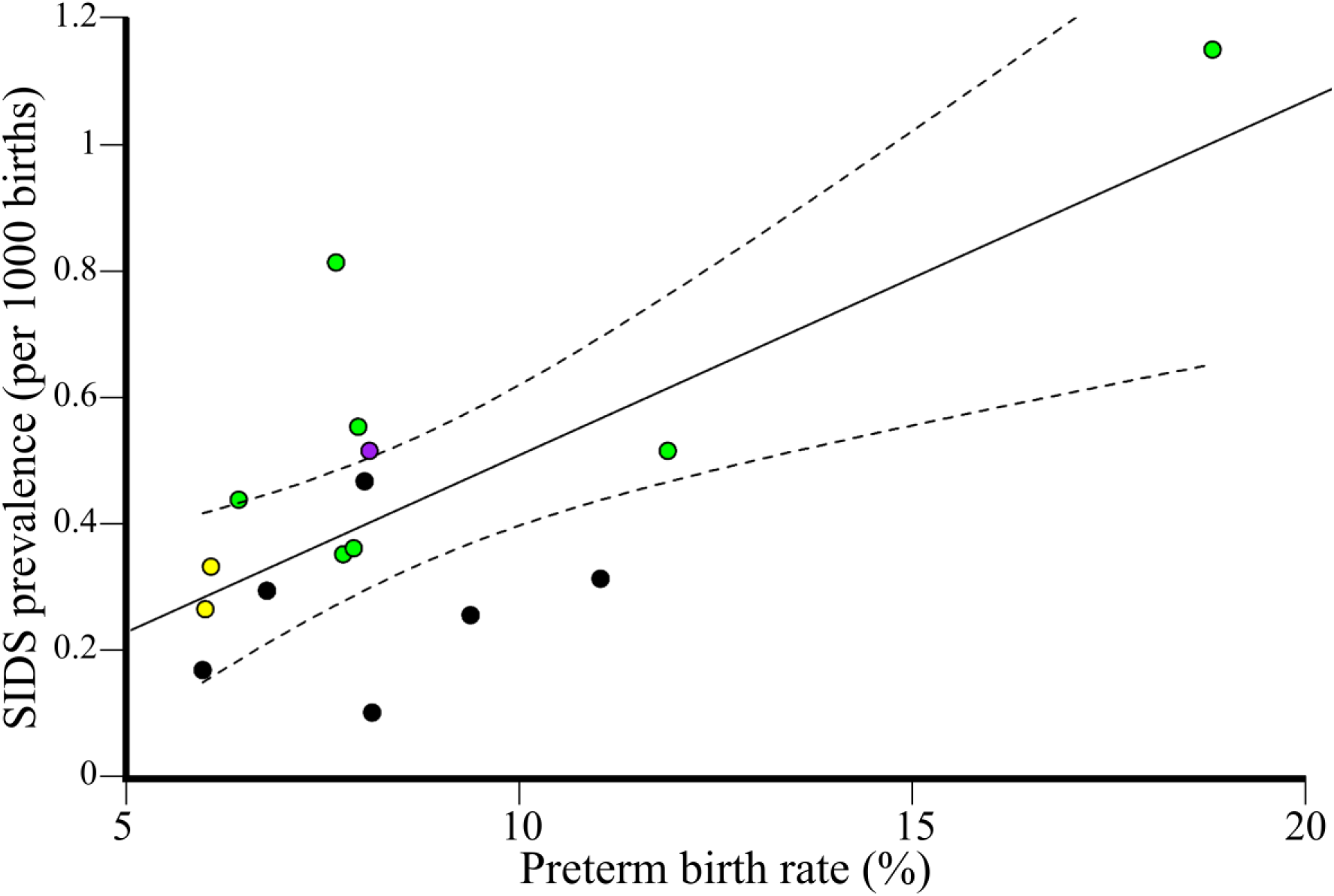
Regression analysis of global prematurity and SIDS prevalence. Data were obtained for 15 states and 16 populations. The 95% confidence intervals of the best fit line are denoted in dashed lines. Color codes are as in Figure 3.

US states also exhibit a significantly positive correlation between SIDS prevalence and prematurity (*N*=50, *r*=0.49, Student’s *t*-test, *p-value*=3*10^−4^, 95% CI: 0.25-0.68) (Figure 7). The slope of this trend indicates that a 10% increase in the prematurity rates is associated with an increase of 1.5 per 1000 SIDS cases. Both prematurity rate and SIDS prevalence are significantly higher in the 16 Southern compared to the 9 Northeastern US states (Kolmogorov-Smirnov test, *p-value*=1.88*10^−5^ and *p-value*=0.004, respectively), suggesting that prematurity is associated with SIDS.

**Figure 7.**
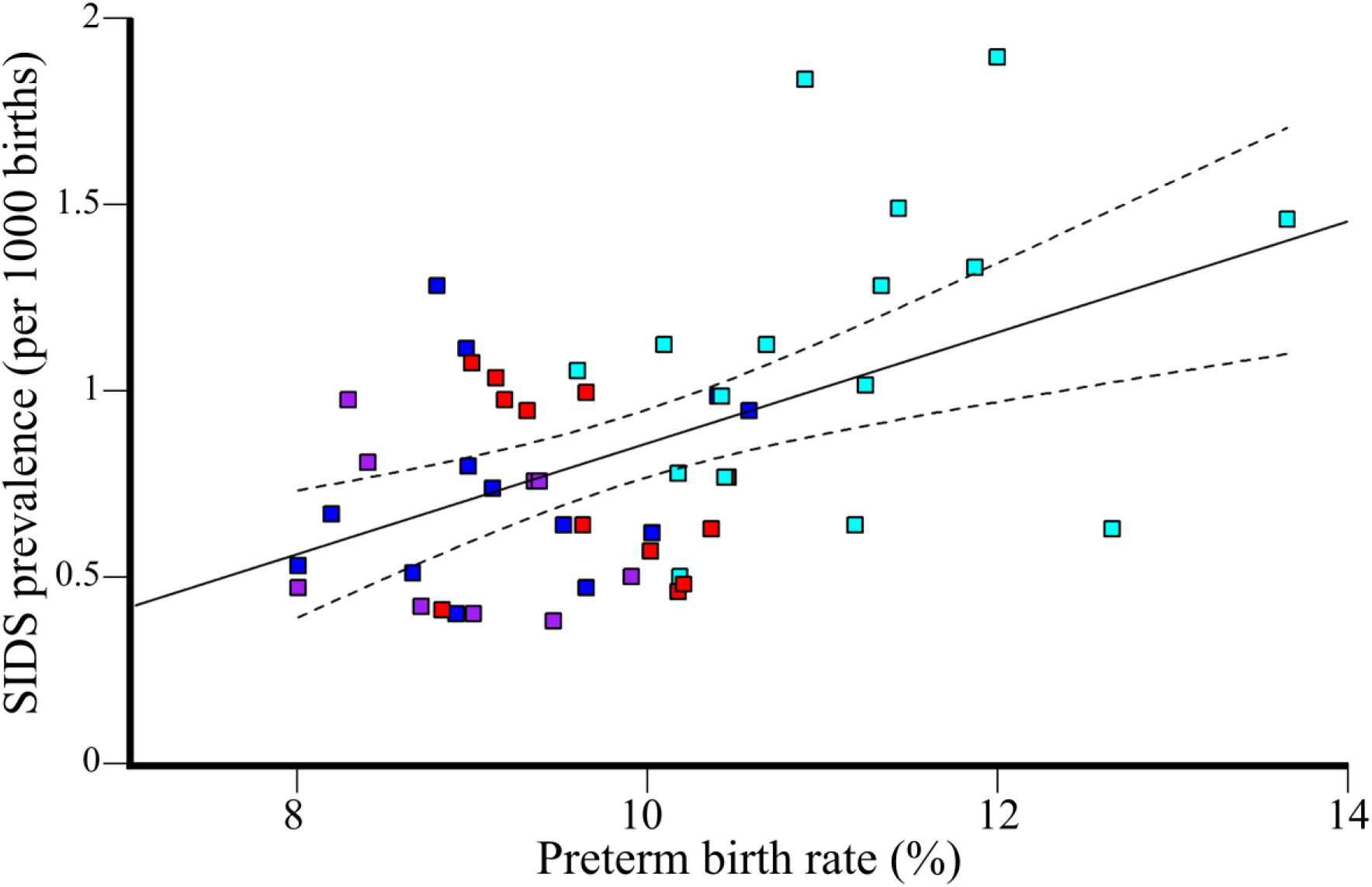
Regression analysis of prematurity and SIDS prevalence in 50 US states. The 95% confidence intervals of the best fit line are denoted in dashed lines. Color codes are as in Figure 2.

Due to the known male bias in preterm births [11], we tested whether prematurity rates explain the gender bias in SIDS in US states. We found insignificant correlation between the prematurity rate and SIDS gender ratio (*N*=32; *r*=0.21, Student’s *t*-test, *p-value*=0.23). Prematurity and MNC rates are not correlated (*N*=39; *r*=0.26, Student’s *t*-test, *p-value*=0.11). In the global dataset, MNC and prematurity were significantly correlated (*N*=16; *r*=0.66, Student’s *t*-test, *p-value*=0.005, 95% CI: 0.24-0.87), suggesting a potential confounder effect.

### Additive effects of various stressors increase the risk of SIDS

To assess the additive effect of MNC and prematurity, we performed likelihood ratio tests considering all possible combinations of the stressors. We found in both the global and US datasets that the combination of MNC and prematurity is significantly or marginally significantly better predictor of SIDS compared to MNC (*p-values_Global_*=0.051, *p-values_US_* 0.018) or prematurity (*p-values_Global_*=0.0002, *p-values_US_* 0.055) alone.

## Discussion

In spite of continuous research and global BTS campaigns, SIDS remains the leading cause of death among infants between birth and 1 year of age [3]. Much of the difficulties in studying SIDS pertain to terminological [43] and methodological issues [44]. SIDS is a diagnosis of exclusion given when the cause of death cannot be determined. Therefore, SIDS can be expected to decrease over time as parental education and pathological diagnoses improve. Indeed, SIDS prevalence has been declining worldwide since the 1980s [45] and has been accommodated by an increase in the prevalence of sudden and unexpected infant deaths (SUIDs) – a diagnosis used to describe the sudden and unexpected death of a baby less than 1 year old in which the cause of death was not obvious before an investigation [46]. Interestingly, SUID prevalence have not improved in the past decade [47], which attests to the limited success of the BTS campaign in actually preventing deaths [45]. Though SIDS prevalence decreases with time as more causes of deaths are becoming known with time, it may also decrease due to the variability in, and confusion about, categorizing deaths [48] or inconsistency between investigators [49]. The causes of death may also intentionally be misrepresented in order to avoid an autopsy due to cultural or religious practices or to avoid a time-consuming investigations [34]. Ontario, for example, eliminated all SIDS-related deaths between 2014 and 2016 by re-categorizing them as “undetermined” deaths [44]. Daunting methodological problems are also prevalent in SIDS studies. The unavailability of proper controls and inability to account for the different life histories of infants beginning *in utero* and their exposure to environmental stressors later in life [e.g., 50] is a major limitation in SIDS studies. Cohort studies are also problematic due to the difficulty of finding suitable controls and accounting for external stressors, which vary widely among countries, cultures, and socio-economic status and can render association studies ambiguous. These methodological difficulties have resulted in over 100 explanations for SIDS that appeared in *Medical Hypotheses* [5] and much confusion between cause and effect. For instance, it has been reported that breastfeeding for a duration of at least two months is associated with a reduced risk of SIDS [51], however, it does not mean that breastfeeding confers protection against SIDS, as an infant’s refusal to feed may be a symptom of other SIDS risk factors, like MNC [52, 53].

The misunderstanding of SIDS is best demonstrated by the popular *triple risk hypothesis* [54], which emphasizes prenatal injury of the brainstem. This prediction was evaluated in a comprehensive SIDS investigation that sequenced the full exons of 64 genes associated with SIDS in 351 infant and young sudden death decedents [55]. Less than 4% of unexpected deaths were associated with a pathogenic genetic variant. Not only has this hypothesis failed to explain the main characteristics of SIDS, but its central argument remains unsupported by the data.

The *allostatic load hypothesis*, initially proposed to explain how stress influences the pathogenesis of diseases [56] and later applied to specific disorders [e.g., 57], proposes that prolonged and repetitive stressful, painful, and traumatic experiences during the peri- and pre-natal developmental periods lead to the accumulation of allostatic load that may be lethal [5]. Thereby, both hypotheses consider external stressors but disagree on the definition of at-risk infants. The *triple risk hypothesis* is more deterministic and implies that, in some cases, SIDS is unavoidable. The *allostatic load hypothesis* considers any infant to be at risk in a direct proportion to their genetic vulnerabilities and the cumulative stressors they have experienced (a wear and tear process) [5].

Here, we tested some of the predictions of the *allostatic load hypothesis* [5]. Due to the aforementioned terminological and methodological problems, we sought to focus on the “low hanging fruits” – the risk-factors that may explain the characteristics that distinguish SIDS from other deaths: MNC and prematurity. Since most of these factors are not recorded during autopsies and thereby cannot be studied retroactively, we carried out an ecological study. We found a positive correlation between SIDS prevalence and common environmental stressors such as neonatal circumcision and prematurity. Together, these stressors were more strongly associated with SIDS then each one separately, suggesting an additive effect, in support of the *allostatic load hypothesis* [5]. The positive correlations between environmental stressors and SIDS are suggestive of the perilous effect that painful and stressful experiences have on infants, particularly vulnerable ones.

### Evaluating the contribution of NMC toward SIDS

It is well-established that male infants are more susceptible to SIDS than females, but the reason is unclear [58]. The genetic explanations for this phenomenon point at the physiological differences for cerebral blood flow, neonatal stress. and various indices of respiratory function in preterm infants [59] and suggest that preterm males need more respiratory support than females [60]. Other explanations proposed that there exists an X-linked *dominant* and protective allele (*p*=1/3) to terminal hypoxia, which leads to a 50% excess in the risk of death for males [61], alternatively there may exist a non-protective X-linked *recessive* allele (*p*=2/3) and a protective dominant corresponding X-linked allele (*q*=1/3) [62]. These explanations assume that the ~0.6 average gender bias in US SIDS cases is biologically meaningful. However, the average gender bias in US SIDS cases is lower than 0.6 (Table S2) and inconsistent among US populations (Table S3). Genetic factors also cannot explain why European countries exhibit different male biases than US states [63, 64].

That SIDS does not have a congenital or genetic origin precludes the existence of major genetic anomalies [65] and highlights the importance of non-genetic factors. When SIDS prevalence differs between various populations that share the same environment, exploring cultural differences can highlight risk factors for SIDS. For instance, the variability in SIDS prevalence (1998–2003) between South Asians (0.2/1000 live births) and White British (0.8/1000) infants who lived in Bradford was explained by the maternal smoking, non-breast feeding, sofa-sharing, and alcohol consumption that were more prevalent in the latter group [66]. In the Netherlands, the higher SIDS prevalence (1996–2000) in Turkish (0.24/1000) and Moroccan (0.28/1000) infants compared to White Dutch ones (0.16/1000) was associated with risk increasing customs unique to each group (e.g., side sleeping and the use of pillows). The dangerous combination of bed-sharing and maternal smoking is a common theme identified by several studies that explored the disparities in SIDS prevalence between different cultures [66, 67]. Yet these risk factors cannot explain the high SIDS mortality in US Whites compared to Europeans [34], low SIDS prevalence among Ibero-American populations [34, 68] compared to US Whites [9], and SIDS male-bias observed among US populations.

We argue that the practice of MNC can explain those differences and showed that large proportions of SIDS and SIDS variation between genders in the US can be explained by the MNC rates but not prematurity. Our results suggest that MNC contributes to the high mortality and gender-bias. That the equivalent practice of female genital mutilation is illegal in a growing number of countries [69] further increases that bias. In addition, females benefit from the protective effect of their sex hormones like estrogen against stressful and painful experience early in gestation [70–72]. We thereby surmise that the gender variation in SIDS is due to the dual legal-biological protection that females enjoy and that eliminating or postponing MNC may decimate the gender bias but not completely eradicate it.

Our finding that MNC is associated with SIDS is not surprising. Circumcision is associated with intra operative and postoperative risks, including bleeding, shock, sepsis, circulatory shock, hemorrhage, pain, and long-term consequences [12–14, 73–76] – all of which contribute toward allostatic load [14, 15] and thereby SIDS through various mechanisms [5]. For instance, circumcision reduces the heart rate [20] and together with the loss of blood there is a danger of reducing the blood volume, blood pressure, and the amount of oxygen reaching the tissues [5, 77]. A reduced blood pressure has been associated with obstructive sleep apnea (OSA), a condition where the walls of the throat relax and narrow during sleep, interrupting normal breathing [77, 78]. Unsurprisingly, SIDS victims experienced significantly more frequent episodes of OSA [79]. Preterm neonates experience over twice the rate of bleeding complications than full-term neonates [80]. MNC-related complications are unavoidable [13, 14, 80–82] and in tandem with the lack of evidence of a meaningful and relevant health benefits to the infant, several countries chose to opt out of the operation [83]. For instance, in 1949, Gairdner’s report [84] that 16 out of 100,000 UK boys under 1-year old died due to circumcision prompted the British government to exclude circumcision coverage from the National Health Service.

Until the late 19^th^ century, Jews were the only group practicing exclusively MNC in Europe [23]. It is thereby of interest to ask whether Jews were familiar with the association between MNC and SIDS. Elhaik [5] already showed that MNC was known to be a deadly practice for over a millennium and prompted the splintering of Reform Judaism from Orthodox Judaism in the nineteenth century. Here, we argue that several Jewish customs associated with MNC reflect the footmarks of SIDS, centuries before it was defined. Jewish ritualistic circumcision, as practiced today, emerged only during the second century AD [85]. It was also around that time that the myth of the baby-killer Lilith, a beautiful, taloned foot demoness [86], became prevalent [87]. Originally one of many Mesopotamian demons, Lilith clawed her way through the demonic hierarchy, extending her influence over time until she became Samael’s (Satan) wife around the 13 ^th^ century [86]. Deceiving Lilith into believing that the newborn was a girl by letting the boy’s hair grow and even dressing him in girl clothes during infancy were the most effective means to avoid her harm. This Middle Age tradition [88] is still being practiced among Orthodox and even secular Jews who avoid cutting a boys’ hair for the first three years. Another ancient tradition is the “Night of Watching,” a ceremony held on the night preceding circumcision to guard the newborn throughout the night against Lilith [89]. In some ceremonies the guests were particularly loud throughout the night to prevent the infant from succumbing to death. Overall, these practices are a testament to Jews’ beliefs that 1) sudden death was and still is highly prevalent; 2) there exists a major male bias in these otherwise random infant deaths; 3) circumcision is associated with sudden deaths; and 4) sudden deaths occur at night – all of which are the hallmarks of SIDS. Unfortunately, there are limited data of the SIDS prevalence in Israel due to religious limitations on conducting autopsies [90]. Interestingly, Israeli health officials reported that, unlike in other countries, Israel saw no reduction in SIDS prevalence following the BTS campaign [91].

### Evaluating the contribution of prematurity toward SIDS

The risk of SIDS among preterm infants remained high and unchanged in the US [25] and is inversely associated with gestational age [24]. For instance, infants born between 24 to 27 weeks were three times more likely to succumb to SIDS than term infants [24]. The risk factors for SIDS are similar in preterm and term infants, except for parity, which is not associated with preterm infants [92]. The lowest SIDS prevalence for preterm infants (<37 weeks) was among Asian/Pacific Islander (1995–1997: 92.8 per 100 000; 2011–2013: 65.2 per 100 000) and Hispanics (1995–1997: 130.6 per 100 000; 2011–2013: 101.7 per 100 000) [93]. Despite of the known male bias in preterm births, we found no association between prematurity and the gender bias in US SIDS cases, suggesting the existence of stronger factors that determine the gender bias in US populations. Our analyses confirmed that prematurity increases the risk for SIDS and that premature circumcised infants are at a higher risk, in agreement with recent findings indicating that preterm neonates suffer from high rate of bleeding complications following MNC [80], immaturity of their cerebrovascular control in the first year of life [94], and neurodevelopmental complications [95, 96], which likely contribute toward the high mortality [24, 25].

### The environmental stressors explain the four main characteristics of SIDS

Our findings explain two out of the four main characteristics of SIDS: male predominance is explained by the prevalence of MNC, a male predominant stressor; the lower SIDS rate among US Hispanics compared to Whites is explained by the low prevalence of MNC in Hispanics compared to Whites [42]. The high prevalence of SIDS cases during the winter or between the second and forth months after birth can be tenuously explained by the accumulation of new stressors, like an increase in respiratory illnesses among household members that are in contact with the infant [97] and the increased sensitivity of infants after their antibody protection weans out [98].

### Implications of our findings

Our findings suggest that MNC, the most common unnecessary surgery in the world, is a major risk-factor for SIDS. Circumcised infants living in a stress-fraught environment, born prematurely, or have an existing genetic predisposition to sudden death would be at the highest risk of SIDS. While the risks of preterm births are well recognized, the debate concerning MNC is polarized between ethical concerns [99] and financial motives [100, 101] clouded by alleged medical benefits, with little awareness of the long-term risks for infants. Although the conclusions of our ecological study should be verified in a cohort study with properly matched infants [102], some recommendation can be implemented immediately at little cost, such as: eliminating neonatal circumcisions when possible, postponing non-medical circumcisions to later ages, informing parents of the risks in MNC, and applying pain management techniques to neonates that experience repetitive pain. MNC data should also be collected and tested in prospective SIDS studies.

### Limitations

This study has significant limitations, many of which are not due to the study design and are common to all SIDS studies. First, as in all ecological studies, correlation is not causation, and causation cannot be inferred from correlation alone. Second, SIDS prevalence data were obtained from 15 countries, which reduced the power of our analyses and may have generated Type I\II errors. Third, pain management techniques practiced in various countries could not be accounted for in our study. Fourth, homogeneity of environmental exposure and diagnosis among the SIDS studies has been assumed, but each may be subject to misclassification, confounding, and biases. Fifth, we assumed the absence of neonatal female circumcision, which is illegal or uncommon in the studied countries. Six, the CDC lists SIDS for all autopsied and nonautopsied cases without distinction. In the case of an interracial parentage, the CDC only reports a single race, usually the one chosen by the mother. Finally, countries measure SIDS in different ways, which can contribute discernibly toward the variation in SIDS prevalence across countries [34]. Changes in the classification of deaths from SIDS to other categories (such as “unknown”) would reduce SIDS prevalence and its association with stressors [103, 104]. Unavailability of same-year data for SIDS and the stressors may also bias their association.

Some of the above-mentioned limitations were addressed by restricting our analyses to countries that perform autopsies and assembling a secondary dataset of US states. Although the age of inclusion for SIDS differs across countries, the difference centers on the inclusion of the first week of life, a time when a meager percentage of SIDS deaths occur [9, 34]. SIDS prevalence and the stressors’ rates do not change dramatically over time [e.g., 9, 34, 105], thus accepting mismatched dates up a few years would likely have small effect on the results. A major difficulty is to find year-matched MNC and SIDS rates globally. We addressed this problem by deriving the low MNC rates from the proportion of Jewish and Muslims populations who tend to remain constant over short period of times and showed that halving or doubling their proportions does not change the results. Stang [106] found that most doctors and obstetricians who perform circumcisions avoid using anesthesia due to the extended time the procedure requires (half-hour) and its potentially negative effects [107–109]. Some of the remaining limitations may be addressed in a carefully constructed cohort studies, but it is likely that other limitations cannot be addressed, in which case our confidence in the associations depends their replicability. For that, we showed that the global and US dataset yield similar patterns and results in agreement with the biological and historical data and in support of the *allostatic load hypothesis* for SIDS [5].

## Conclusions

SIDS is a diagnosis with a multifactorial underlying etiology. The *allostatic load hypothesis* [5] explains the main characteristics of SIDS (male predominance, different rates among US group, prevalence peak between 2 and 4 months, and seasonal variation) in the prolonged and repetitive stressful, painful, and traumatic stimuli that may begin prenatally, tax neonatal regulatory systems, and increase the risk of SIDS. Our ecological analyses support an association between MNC, prematurity, and SIDS and the additive effects of MNC and prematurity toward SIDS. Mitigating these and other stressors may reduce the prevalence of SIDS. Our data and code can be used to evaluate associations with other environmental factors. Future cohort studies should consider the existence of these stressors, genetic vulnerabilities, and life history.

## List of abbreviations

(SIDS): Sudden Infant Death Syndrome
(MNC): Male Neonatal Circumcision
(BTS): Back To Sleep
(OSA): Obstructive Sleep Apnea

## Ethics approval and consent to participate

Not applicable

## Consent for publication

Not applicable

## Availability of data and material

All the data and *R* scripts to generate our figures are available via GitHub.

## Acknowledgment

Not applicable

